# Remote-control meiotic drive of sex chromosomes

**DOI:** 10.1101/2024.11.18.624228

**Authors:** Naomi L. Greenberg, Manus M. Patten, Martijn A. Schenkel

## Abstract

Some selfish genetic elements drive at meiosis to achieve transmission distortion, breaking the rules of Mendelian segregation to enhance their own evolutionary success. It has been shown that enhancers of drive must act in *cis* in order to gain the selfish benefit of drive and that suppressors of drive will be selected at unlinked loci. Here, we model the evolution of an autosomal *trans*-acting gene (*Distorter*) that causes the Y-chromosome (or even 0-chromosome) to drive without driving itself, a phenomenon we call “remote-control meiotic drive”. We show that such a gene may spread in the population when linked to a second locus, *Assister*, whose alleles are transmitted at different frequencies through sperm as compared to eggs, for which we consider various scenarios, e.g. sexually antagonistic selection or sex-limited drive. Depending on the mechanistic details of sex-chromosome drive, *Distorter*’s spread can additionally facilitate transitions between XY and X0 sex determination. Our results provide a proof of principle that stretches the current understanding of segregation modifier and sex allocation theory. Moreover, we identify alternative evolutionary trajectories that could also lead to remote control drive, and discuss its potential applications in developing synthetic sex-ratio-distorting elements for use in e.g. pest management.

## Introduction

Mendel’s Law of Segregation states that in a diploid individual, the two alleles at a locus segregate to an equal (i.e., 50/50) proportion of the gametes (Mendel, 1865). The fair transmission of genetic material can be distorted in myriad ways (Burt & Trivers, 2006; Patten *et al*., 2023), such as by true meiotic drivers (Clark & Akera, 2021) and, to similar effect, post-segregational killers (Bravo Núñez *et al*., 2018), which often fall under the moniker of “meiotic drive” themselves. The former achieve their advantage in asymmetric meiosis by segregating preferentially to the egg pole during meiosis and avoiding the polar body (Sandler & Novitski, 1957). The latter target their allelic counterpart just after meiosis, e.g. by preventing the formation of functional gametes that lack the transmission-distorting gene (Lindholm *et al*., 2016; Price *et al*., 2020). Consequently, the gametes produced by heterozygotes do not feature equal proportions of both alleles; instead the transmission-distorting allele becomes overrepresented, and the non-distorting allele underrepresented. Meiotic drivers typically harm the fitness of their bearers, often by causing reduced fertility (Lyttle, 1991; Larracuente & Presgraves, 2012). In the case of sex-chromosome drive, additional harm comes from overproducing the common sex (Lyttle, 1991). In either case, their ability to cheat the Law of Segregation offsets this cost to themselves, enabling them to spread and persist in the population.

In males, meiotic drivers may achieve their higher-than-Mendelian transmission using two mechanisms, both of which entail a supergene that consists of two distinct loci. First, the toxin-antidote system (Burga *et al*., 2020) features a toxin that is expressed during or prior to meiosis, so that all resulting gametes receive a dose. In gametes that bear the toxin-antidote supergene, the antidote is expressed post-meiosis to prevent the harmful effects of the toxin and rescue the supergene-bearing gametes. The second system is the killer-responder system, which produces a killer element that targets a responder locus (Meiklejohn & Tao, 2010). In this case, the killer allele is linked to an insensitive variant of the locus that is targeted by the killer, whereas the non-killer allele is assumed to be linked to a sensitive variant and is therefore eliminated. Gametes that lack the supergene (or more specifically, the antidote allele or the insensitive responder allele) are eliminated, so that the gamete pool produced consists predominantly of gametes bearing the segregation-distorting complex at the supergene.

A key feature of both mechanisms is that the targeted genetic element is the chromosomal complement of the driver, meaning segregation distortion acts to remove these from the gamete pool to enhance its own odds of being transmitted. Theoretical models have explored how this dynamic affects the potential for genes that modify the severity of segregation distortion. According to modifier theory, genetic modifiers that arise linked to the segregation-distorting complex would typically act to enhance the degree of segregation distortion, whereas modifiers linked to the targeted gene complex would be expected to diminish its effect (Prout *et al*., 1973; Feldman & Otto, 1991). Effectively, this constitutes the formation of antagonistic, multigenic collaborations to cheat Mendelian transmission or enforce it, respectively. Unlinked modifiers, i.e. those occurring on other linkage groups, are also typically thought to favor diminished segregation distortion owing to the deleterious effects segregation distorters have on their bearer’s fitness (Eshel, 1985; Veller, 2022). These dynamics underlie the so-called “parliament of genes” effect which seeks to explain why so few genes succeed in being selfish (Leigh, 1971). Only the select few genes linked to a selfish genetic element like a segregation distorter are selected to enhance it, but the remainder of the genome is selected to suppress it. Through sheer numbers, the latter typically ends up as the winner.

As a variation on the above, we could consider a segregation distorter that targets not its own chromosomal complement, but rather one of another linkage pair. One hypothetical example of this is an autosomal killer gene that targets a responder gene on one of the sex chromosomes; this might also be considered a modifier gene that affects the segregation of the sex chromosome pair (Maffi & Jayakar, 1981). Such a gene would normally be counter-selected for two reasons. The first is that by biasing the sex ratio among its bearer’s offspring, it ultimately ends up in the overrepresented sex (Lyttle, 1991; Jaenike, 2001). The second is that the segregation distorter would target gametes containing copies of itself half of the time (i.e., “suicide combinations”; Hartl, 1974). Such *trans*-acting or “remote-control” meiotic drive has been largely ignored, as the lack of benefit obtained from eliminating the competition and potential additional costs to organismal fitness of such a segregation distorter are thought to render it unlikely to evolve and spread. In this manuscript, we identify a route by which such a mechanism may overcome these costs and lead to the evolution of autosomal genes for sex chromosome meiotic drive.

Drive may be found more commonly on sex chromosomes than autosomes for two reasons, one owing to genetics and the other to ascertainment bias. First, the X- and Y-chromosomes do not typically recombine for much of their length, which prevents the separation of the alleles that bring about segregation distortion when found in tandem (Hartl, 1974; Haig & Grafen, 1991). Repeated cycles of drive and suppression, which may be common on the X- and Y-chromosomes for this reason, may lead to the evolution of many cryptic drivers on the sex chromosomes (Ellison & Bachtrog, 2019; Bachtrog, 2020). Second, before suppressors of sex-chromosome drive evolve, they are more readily ascertained than autosomal segregation distorters, as skewed transmission of the sex chromosomes brings about a conspicuous shift in the offspring sex ratio (Hamilton, 1967; Jaenike, 2001, 2008). Further, this sex ratio effect can maintain drivers in a polymorphic state owing to the frequency dependence of fitness. The sex chromosomes may therefore be enriched for both fixed cryptic drivers as well as polymorphic drivers.

Sex ratio biases brought about by drive can promote evolutionary transitions between sex determination mechanisms (Kozielska *et al*., 2010). For example, a driving X-chromosome may lead to female-biased sex ratios, promoting the evolution of a male-determining gene on an autosome, converting it into a proto-Y-chromosome. Autosomal driving elements may also promote the origin of sex chromosomes (Úbeda *et al*., 2014). Owing to differences between oogenesis and spermatogenesis, most meiotic drivers exhibit sex-specific (or sex-biased) segregation distortion, so that one sex may transmit it at a Mendelian rate, whereas in the other it distorts segregation to achieve higher transmission. If a locus linked to this sex-specific driver evolves a novel, sex-determining mutation, this confines the driving allele to the sex in which it achieves segregation distortion more frequently. As this supergene spreads in the population, the segregation-distorting effect causes sex-ratio biases (similar to Kozielska *et al*. (2010)), leading to a series of selectively favored mutations that eventually yield a set of sex-determining chromosomes with normal, Mendelian segregation.

The origin of a sex-determining gene next to a segregation-distorting gene effectively restricts the distorter to one sex, forming a synergistically-acting supergene. We posit here that achieving this effect does not explicitly require a sex-determination gene, but rather that biasing transmission to one sex more generally enables this to occur. That is, any gene that skews the sex ratio of its bearers’ offspring would produce a similar effect. Ordinarily, a sex-ratio-biasing autosomal gene would be harmful to its bearers, and hence be purged (per Fisher (1930), but see West (2009)). But a sex-ratio-biasing gene paired with a sex-specific meiotic driver establishes a synergism that offsets such costs and might therefore be able to spread and persist.

Here, we model the evolution of an autosomal, *trans*-acting segregation distorter (*Distorter*) that biases the transmission of the sex chromosomes. We consider two potential proximate mechanisms by which it achieves this effect, which we dub X-shredding and X-conversion, further explained below. Linked to this *trans*-acting segregation distorter is a secondary locus (*Assister*), which may represent: (1) a locus under sexually antagonistic selection, where different alleles are favored in females as compared to males (van Doorn, 2009; Schenkel *et al*., 2018) or (2) a sex-specific meiotic driver, an analog to the situation modeled by Úbeda *et al*. (2014). In the Supplement we consider two other scenarios, one in which the *Assister* locus is parentally antagonistic, wherein an allele’s fitness depends on its parental origin (Patten *et al*., 2013; Haig *et al*., 2014), and a second in which the *Assister* locus simply causes co-segregation of the autosome and the Y-chromosome. The underlying rationale for these scenarios is that in all cases, the *trans*-acting segregation-distorting gene and the linked gene have the same optimal destination—namely, a male offspring (Gardner & Úbeda, 2017). In addition, these phenomena have previously been shown capable of spurring transitions between sex-determining mechanisms and of affecting sex ratio adjustment (van Doorn & Kirkpatrick, 2007, 2010; Blackburn *et al*., 2010; Kozielska *et al*., 2010; Úbeda *et al*., 2014; Muralidhar & Veller, 2018; Schenkel, 2024). We use our model to determine the conditions under which such a gene can spread in the population, what frequency it reaches at equilibrium, and what level of sex-ratio distortion is achieved once equilibrium is attained by causing increased transmission of the Y-chromosome over the X-chromosome. We also consider whether it can spur a transition between genetic sex-determination mechanisms, where this interference results in the formation of 0-chromosomes, whose spread in the population leads to a loss of the Y-chromosome. Through this, we show that autosomal genes may interfere with the segregation of the sex chromosomes to promote their own spread in the population. This results in the sex chromosomes appearing to drive through no effort of their own—i.e., through remote control. We discuss our results in light of previous modifier theory of segregation distortion, sex allocation theory, definitional issues of gene drives, and in the context of potential applications.

## Methods

Our model assumes an infinite population with XY sex determination (females XX, males XY), with an initially equal sex ratio. The X- and Y-chromosomes determine sex and do not carry any other genes relevant to our model. Terms such as ‘female’ and ‘male’, and similarly ‘XY’ versus ‘ZW’ can be exchanged to consider cases with female heterogamety, rather than male heterogamety. In addition to the XY sex chromosomes, individuals carry an autosome carrying two loci, dubbed *Distorter* (with alleles *D*_1_ and *D*_2_) and *Assister* (with alleles *A*_1_ and *A*_2_); recombination between *Distorter* and *Assister* occurs at a rate *r* in both sexes (Figure 1A). The *D*_1_ allele makes for equal segregation of the sex chromosomes, whereas the *D*_2_ allele interferes with the normal (i.e., Mendelian) segregation of the XY sex chromosomes in males. Further details on *D*_*2*_’s mechanism are provided under “*Trans*-segregation distortion by *Distorter*” below.

**Figure 1:**
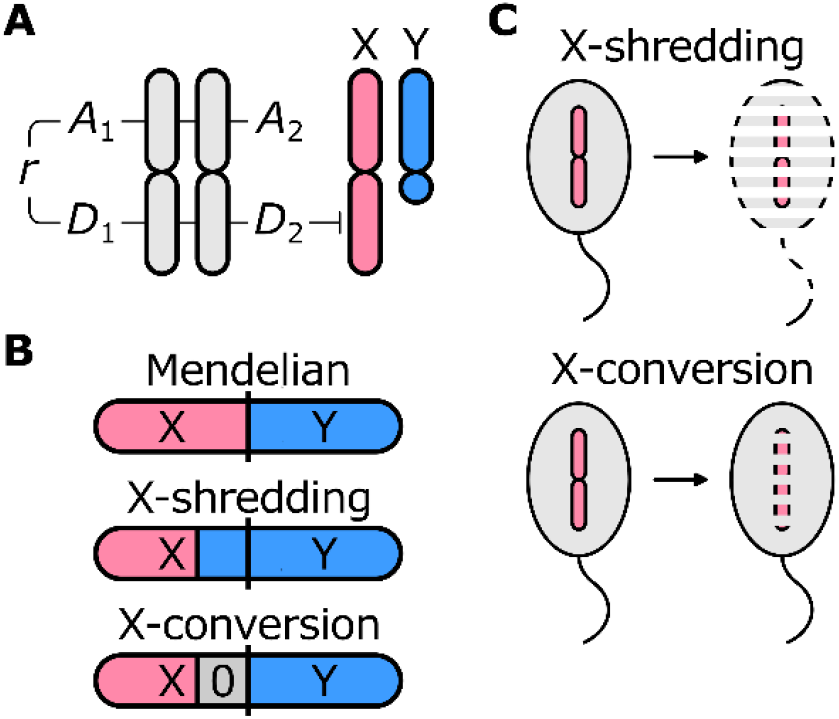
Model overview. (A) Our model incorporates two linked autosomal loci, *Assister* (with alleles *A*_1_ and *A*_2_) and *Distorter* (with alleles *D*_1_ and *D*_2_); recombination between *Assister* and *Distorter* occurs at a rate *r* in both sexes. *Assister* may represent either a sexually antagonistic locus (*A*_1_ beneficial in females/detrimental in males, *A*_2_ detrimental in females/beneficial in males) or a locus experiencing *cis*-segregation distortion (*D*_2_ overtransmitted in *D*_1_/*D*_2_ heterozygotes). The *D*_2_ allele interferes with the Mendelian transmission of the X-, Y-, and/or 0-chromosomes during spermatogenesis to achieve *trans*-segregation distortion.

For the *Assister* locus, we focus in the main text on two selective scenarios, sexually antagonistic selection and male-limited *cis*-segregation distortion, and examine the cases of parentally antagonistic selection and of co-segregation in the Supplementary Material. More details of the two schemes we examine in the main text are provided in “Sexually antagonistic selection on *Assister*” and “*Cis*-segregation distortion by *Assister*”, respectively. The crux of each of these selective scenarios is that the *A*_2_ allele experiences higher long-term fitness by segregating to males, either because: (1) this is the sex in which it exhibits higher-than-Mendelian transmission (segregation distortion); or (2) it enhances the sex of its bearer (sexual antagonism); or (3) it comes to be transmitted in a manner that enhances its fitness in the subsequent generation (parental antagonism). This favors the buildup of linkage disequilibrium between *Assister* and *Distorter*, as well as statistical linkage between this pair of loci and the Y-chromosome. Consequently, the *trans*-segregation distorting allele *D*_2_ can spread via its association with the *A*_2_ allele and its overrepresentation in males. We measure linkage disequilibrium in sperm to track these non-random associations, using the following equations (Rydzewski *et al*., 2016):

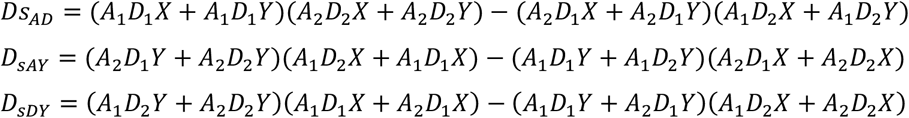

And for the X-conversion model, which additionally includes 0-chromosomes as a third allele at the sex chromosomes:

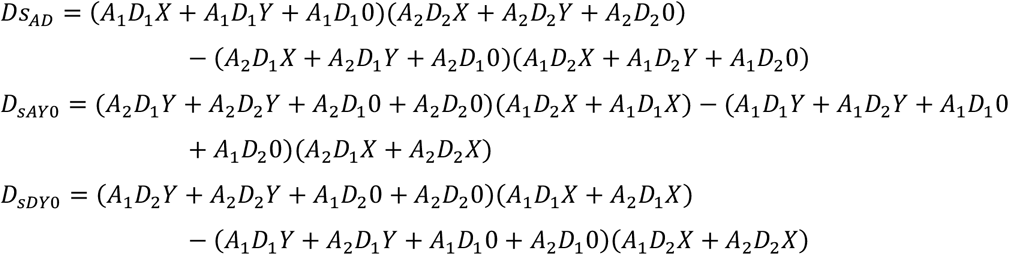

The overall life cycle is as follows: mature males and females undergo meiosis to produce their respective gamete pools; during this time, segregation distortion takes place; gametes are then united at random; the resulting juvenile offspring undergo viability selection during maturation; and upon reaching maturity they replace their parents as the reproductive population so that generations are non-overlapping. At this point, the cycle resets. Our population is initially fixed for the *D*_1_ allele, and with *A*_1_ and *A*_2_ at or near their equilibrium as achieved by 500 “burn-in” generations of simulation. We then mutate a small (*p*=0.001) proportion of *D*_1_ alleles to *D*_2_ alleles on each genetic background in Y-bearing sperm simultaneously (proportional to their frequency in the population at the time) to prevent any advantage *D*_2_ might experience from arising under linkage disequilibrium. The eventual results did not differ when we introduced *D*_2_ on alternative genetic backgrounds instead. We then continue the simulation procedure for 1,000 generations before we evaluate the terminal frequency of *D*_2_.

In our simulations, we vary the parameter values for the selective effects of the *Assister* and *Distorter* loci, as well as their capacity to distort allelic segregation during meiosis (where applicable); details on these are provided under “Results”. We focus here in the main text primarily on the impact of the strength of selection on *Assister* (for sexually antagonistic selection) or its capacity to achieve higher-than-Mendelian transmission (for *cis*-segregation distortion), as well as the strength of *trans*-segregation distortion achieved by *Distorter*. We consider the role of recombination between the *Assister* and the *Distorter* loci and direct selection against *Distorter* more fully in the Supplement.

An overview of all model variables, their descriptions, and standardized values is included in Supplementary Table S1.

### Sexually antagonistic selection on *Assister*

Under sexually antagonistic selection, the *A*_1_ and *A*_2_ alleles differently affect the fitness of female versus male bearers. We assume that *A*_1_ is beneficial to female fitness but costly to male fitness, and vice versa for *A*_2_. The partial fitness score 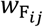 of a female with a genotype *A*_*i*_*A*_*j*_ is given by:

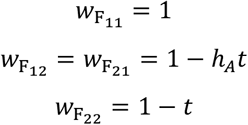

The partial fitness score 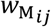 of a male with a *A*_*i*_*A*_*j*_ genotype is given by:

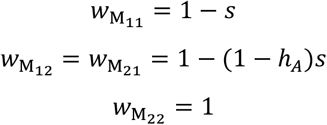

Here, *t* and *s* indicate the homozygous fitness cost of *A*_2_ in females and *A*_1_ in males, respectively, and *h*_*A*_ ∈ [0,1] is the dominance of *A*_2_ in heterozygotes.

### *Cis*-segregation distortion by *Assister*

As a segregation-distorting gene, the *A*_2_ allele of the *Assister* locus is transmitted at a higher-than-Mendelian rate in heterozygotes. Specifically, we assume that an *A*_2_*A*_1_ (or *A*_1_*A*_2_) heterozygote transmits the *A*_2_ allele at a rate of 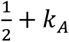 and the *A*_1_ allele at a rate 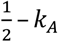 where 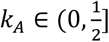 indicates the strength of *cis*-segregation distortion by *A*_2_ and *k*_*A*_ = 0 represents Mendelian segregation. We assume that the *A*_2_ allele negatively affects its bearer’s fitness in females and males alike. The partial fitness score *w*_*ij*_ of an individual with genotype *A*_*i*_*A*_*j*_ is given by:

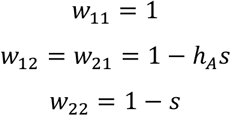

Here, *s* ∈ [0,1] represents the fitness effect of the *A*_2_ allele in homozygotes, *h*_*A*_ ∈ [0,1] its dominance in heterozygotes. We chose this fitness scheme for *Assister* to reflect some well-known autosomal drivers (e.g., the *t*-haplotype in mouse and *SD* in *Drosophila*, which have linked recessive lethality/sterility mutations) and also because it maintains polymorphism at the *Assister* locus, which is essential for our model.

### *Trans*-segregation distortion by *Distorter*

The *D*_2_ allele of the *Distorter* locus acts to distort the segregation rate of the sex chromosomes. We assume that *D*_2_ acts only in males, i.e. the heterogametic sex, as a dominant *trans*-acting segregation distorter, such that *D*_2_*D*_1_ heterozygotes and *D*_2_*D*_2_ homozygotes transmit their sex chromosomes at equally distorted frequencies. We consider two mechanisms of *trans*-segregation distortion, denoted X-shredding and X-conversion, respectively. X-shredding represents a skew in the transmission of X-versus Y-chromosomes to future offspring, where the Y-chromosome is transmitted at a rate 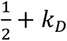 and the X chromosome at a rate 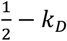, where 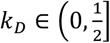; this is achieved by the killing off of some fraction of X-bearing sperm. Under X-conversion, *D*_2_ acts to eliminate the X-chromosome from a certain number of X-bearing sperm to convert them into 0-bearing gametes that, when fused with an X-bearing egg yield a viable, fertile X0 male (note that this assumes that maleness is determined by lack of a second X-chromosome, rather than some dominant male-determining gene on the Y-chromosome, in which case X0 individuals would develop as females). Consequently, the Y-chromosome is transmitted at a Mendelian rate of 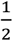, the X-chromosome is transmitted at a rate 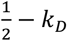, and the remaining *k*_*D*_ represents the proportion of sperm that have been converted from being X-bearing to 0-bearing. In X0 males, the 0-chromosome would be transmitted at a rate 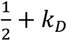and the X-chromosome at a rate 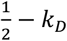. We assume that lacking the Y-chromosome comes at some fitness cost *v* ∈ (0,1] to males, so XY males have a relative fitness score of 1 and X0 males have a fitness of 1 − *v*. Potentially, Y-chromosome-bearing males may also experience reduced fitness owing to the accumulation of mutations on the Y-chromosome, resulting in an effective fitness benefit for X0 males (*v* < 0); we consider this scenario in the Discussion.

We assume that the *D*_2_ allele is harmful to male fertility through its sperm-killing or X-conversion efforts. The resulting partial fitness score *w*_*ij*_ of a male with a *D*_*i*_ *D*_*j*_ genotype is given by:

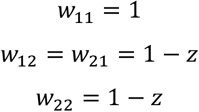

Here, *z* ∈ [0,1] measures the loss of total fertility, and we assume that the *Distorter* locus is neutral in females. Finally, we assume that the partial fitness effects of the *Assister* and *Distorter* loci are multiplicative; Supplementary Table S2 provides an overview of the fitnesses of all potential genotypes in our population.

### Data analysis and visualization

Simulations, data analysis, and visualization were carried out using R (R Development Core Team, 2023; v.4.4.1) and RStudio (RStudio Team, 2023; v.2024.09.0+375) using the ‘tidyverse’ and ‘viridis’ packages (Garnier, 2018; Wickham *et al*., 2019).

## Results

### Sexually antagonistic selection on *Assister* enables the spread of *Distorter*

We first explored whether an autosomal *trans*-acting segregation distorter (i.e., the *D*_2_ allele of the *Distorter* locus) of the Y-chromosome could spread in the population when linked to a sexually antagonistic locus (the *Assister* locus) under rather favorable conditions. With strong sexually antagonistic selection (*s, t* = 0.5), a high level of *trans*-acting segregation distortion (*k*_*D*_ = 0.25) but low fitness costs thereof (*z* = 0.001), and tight linkage between *Assister* and *Distorter* (*r* = 0.001), we found that *D*_2_ can invade and reach a frequency of ∼0.13, enhancing Y-chromosome transmission so that it reaches a frequency – and hence a population sex ratio – of 0.57 (Figure 2A). Meanwhile, the *Assister* locus remains polymorphic, with *A*_2_’s frequency changing only slightly from 0.58 at its initial equilibrium to a frequency of 0.55 after 500 generations (Figure 2A).

**Figure 2.**
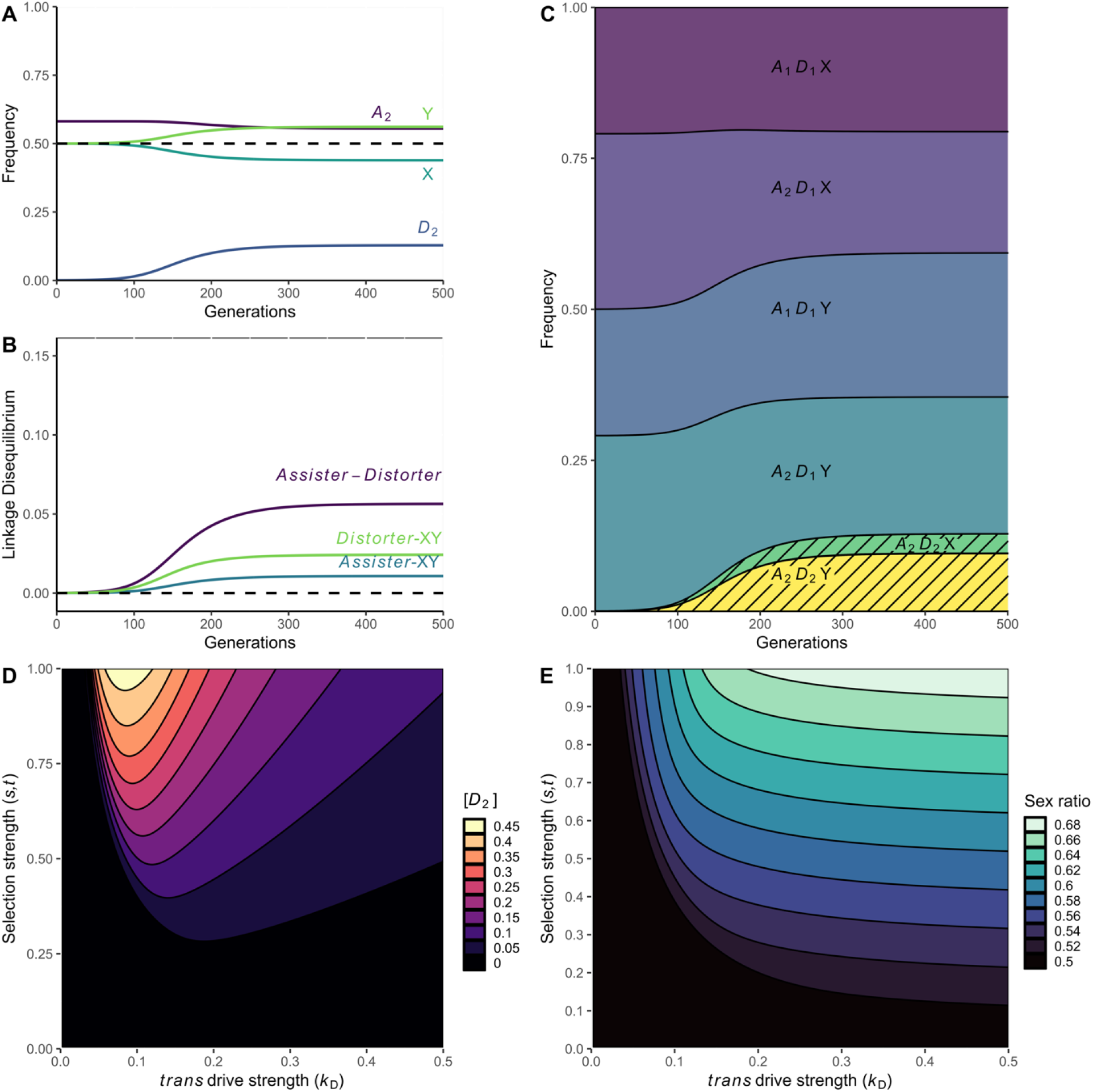
Dynamics and scope of Y-chromosome drive via the *trans*-acting *Distorter*. Here, *Distorter* is linked to a sexually antagonistic *Assister* locus and causes drive via the X-shredding mechanism. (A) Changes in the frequency of the *A*_2_ and *D*_2_ alleles and the X- and Y-chromosomes in males across 500 generations. (B) Changes in the linkage disequilibrium of *Assister-Distorter, Assister-*XY, and *Distorter-*XY across 500 generations. (C) Haplotype frequencies over 500 generations. *A*_1_*D*_2_X and *A*_1_*D*_2_Y have near-zero frequencies. The shaded area represents the frequency of *D*_2_ and primarily comprises the haplotypes *A*_2_*D*_2_Y and *A*_2_*D*_2_X. (D) *D*_2_ equilibrium frequency after 1,000 generations. (E) Population sex ratio after 1,000 generations. Parameter values (if not otherwise specified): *k*_*D*_ = 0.25, *s, t* = 0.5, *h*_*A*_ = 0.5, *z* = 0.001, *r* = 0.001.

We simultaneously tracked the linkage between *Assister, Distorter*, and the XY chromosomes (Figure 2B). These three loci come into close statistical linkage and are inherited non-independently—effectively, as a group—which creates the following reinforcing benefits: *D*_2_ to benefit from the male-beneficial fitness effects of *A*_2_; the Y-chromosome to benefit from its increased transmission thanks to *D*_2_; and *A*_2_ to benefit from being assigned to its fitness-enhancing state (i.e., males) thanks to a stronger association with the Y chromosome.

To better characterize the conditions that favor the spread of *D*_2_, we fully explored the parameter space of sexual antagonism (*s* and *t*) at the *Assister* locus and the strength of drive (*k*) of the *D*_2_ allele. We found that *D*_2_ can spread under most parameter value combinations considered here (Figure 2D), though it achieves the highest equilibrium frequency when *trans* drive is relatively weak (*k*_*D*_ ≈ 0.1) and sexually antagonistic selection is strong (*s, t* ≅ 1). The presence of *D*_2_ in the population leads to biased sex ratios, which are highest for high levels of *trans* drive and strong sexually antagonistic selection (Figure 2E). When sexually antagonistic selection is relatively weak, we see that higher *trans*-segregation distortion does not lead to concomitantly higher frequencies of *D*_2_. Under these conditions, the cost to *D*_2_ of assigning itself to the common sex outstrip the benefits linkage to *A*_2_.

We also investigated the constraints that higher recombination rates and higher *D*_2_ fertility costs place on the spread of *D*_2_. We found that higher recombination (*r* = 0.01 or *r* = 0.1) reduced the opportunity for male-biased sex ratios, likely by reducing the necessary *Assister-Distorter-*XY linkage disequilibrium; similarly, a higher *D*_2_ fertility cost reduced its success (Supplementary Figure S1).

### *Cis*- and *trans*-segregation distortion can work in tandem to achieve sex ratio biases

We then explored whether a male-limited, *cis*-acting segregation distorter at the *Assister* locus could similarly favor the spread of *D*_2_. We found that for a large range of values of *cis* drive strength (*k*_*A*_) and *trans* drive strength (*k*_*D*_), *D*_2_ spreads and the population skews male. Similar to the sexual antagonism model above, the Y-chromosome as well as *D*_2_ increase in frequency, whereas *A*_2_’s frequency remains relatively stable (Figure 3A). The linkage disequilibrium between *Assister* and *Distorter* is strong, as associations between the Y-chromosome and both *D*_2_ and *A*_2_ develop (Figure 3B). *D*_2_ occurs primarily in combination with *A*_2_ (Figure 3C). We investigated the full range of drive strengths (*k*_*A*_ ∈ [0, 0.5], *k*_*D*_ ∈ [0, 0.5]) at both the *Distorter* locus and the *Assister* locus (Figures 3D, 3E). We found that most of the parameter space permits invasion of *D*_2_, and some parameter combinations enable high frequencies of *D*_2_ and a high sex-ratio skew. With freer recombination between *Assister* and *Distorter* and with elevated fitness costs of *D*_2_, the opportunity for *D*_2_’s spread diminishes, and the resulting sex-ratio skew is less extreme (Supplementary Figure 2).

**Figure 3.**
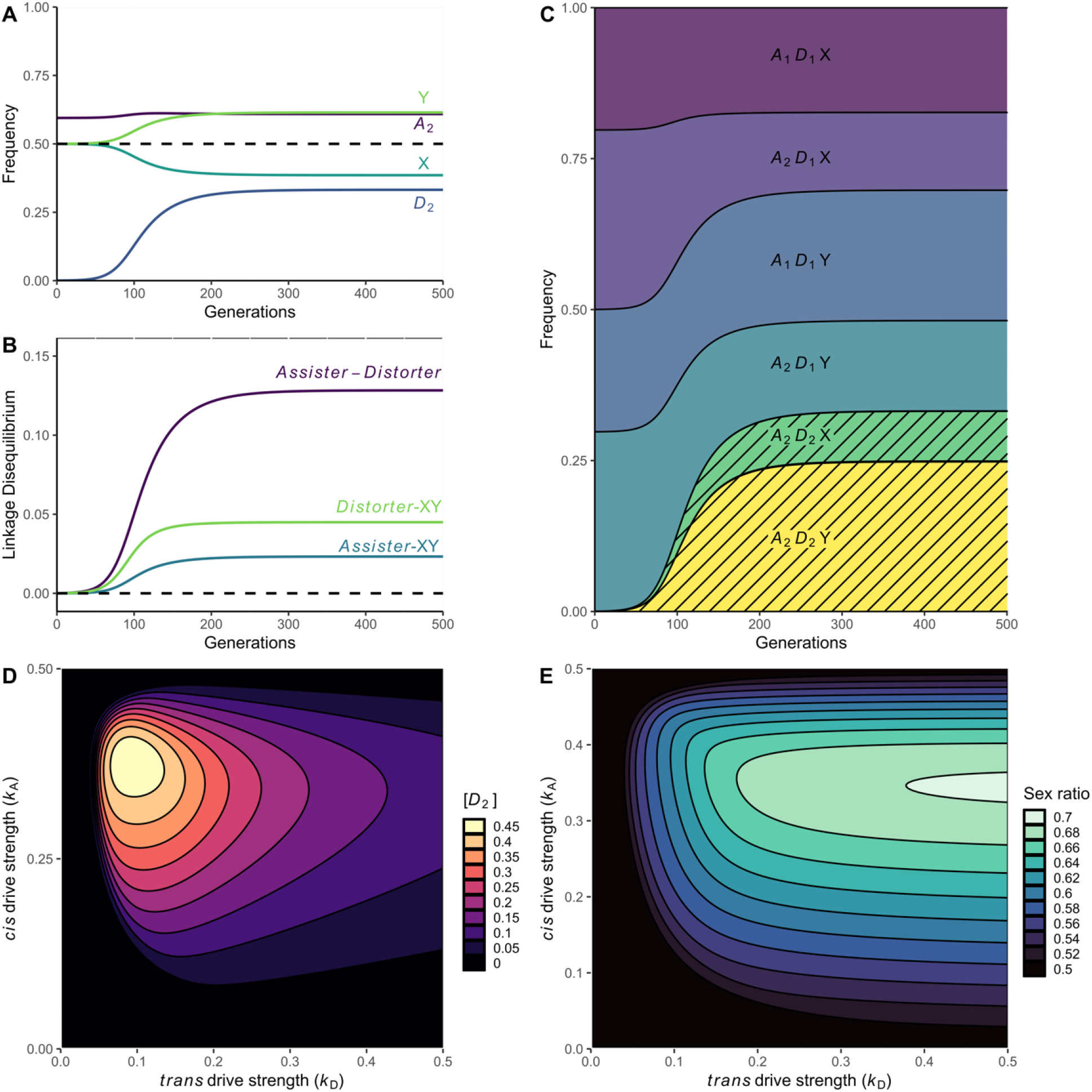
Dynamics and scope of Y-chromosome drive via the *trans*-acting *Distorter* causing X-shredding, linked to a male-limited meiotic driver *Assister*. (A) Changes in the allele frequency of *A*_2_, *D*_2_, and the X- and Y-chromosome in males across 500 generations. (B) Changes in the linkage disequilibrium of *Assister-Distorter, Assister-*XY, and *Distorter-*XY across 500 generations. (C) Haplotype frequencies over 500 generations. *A*_1_*D*_2_X and *A*_1_*D*_2_Y have near-zero frequencies. The shaded area represents the frequency of *D*_2_ and primarily comprises the haplotypes *A*_2_*D*_2_Y and *A*_2_*D*_2_X. (D) *D*_2_ equilibrium frequency after 1,000 generations. (E) Population sex ratio after 1,000 generations. Parameter values (if not otherwise specified): *k*_*D*_ = 0.25, *k*_*A*_ = 0.25, *h*_*A*_ = 0.0, *z* = 0.01, *r* = 0.001, *s* = 0.5.

### X-conversion leads to the emergence of X0 sex determination through loss of the Y-chromosome

In the previous sections, we assumed that *D*_2_ distorts the segregation of the X- and Y-chromosomes by the selective killing of X-bearing sperm. An alternative mechanism by which *D*_2_ might generate a male-biased sex ratio among its bearer’s offspring is to selectively eliminate the X-chromosome from sperm to convert them to 0-bearing sperm, yielding X0 sons instead of XY sons (note however that this assumes that sex determination occurs through some dosage effect of the X-chromosomes as in *Drosophila* fruit flies (Pomiankowski *et al*., 2004); if maleness instead is conferred by a Y-chromosomal dominant male-determiner, an X0 genotype would be expected to lead to femaleness (Graves, 2006), constraining the spread of *D*_2_). This mechanism might not harm male fertility as severely as the X-shredding mechanism, and so is poised to spread more easily. This form of *trans*-segregation distortion also provides a mechanism for an XY sex determination system to transition to an X0 sex determination system. We find that when the *D*_2_ allele causes X-conversion, the resulting 0-chromosomes accumulate and become the predominant male-determining factor. This holds true when *Assister* experiences sexual antagonism (Figure 4), as well as when it exhibits male-limited *cis* drive (Figure 5). In both cases, we find that the 0-chromosome can entirely replace the Y-chromosome. There is only a small range of parameter space in which the Y-chromosome escapes loss and is maintained alongside the 0 chromosome in the population (Supplementary Figure S5).

**Figure 4.**
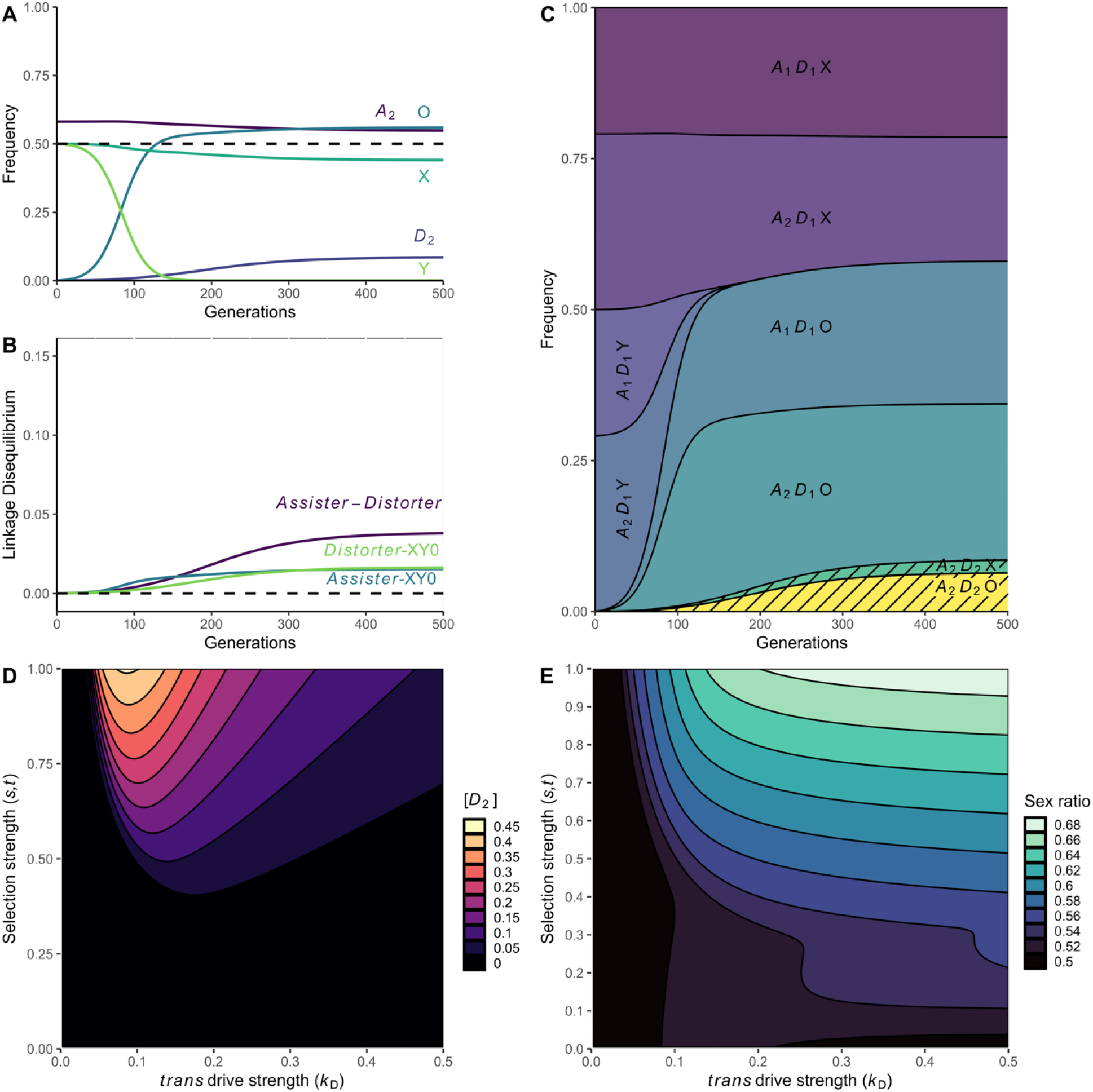
Dynamics and scope of 0-chromosome drive via the *trans*-acting *Distorter* causing X-conversion, linked to a sexually-antagonistic *Assister*. (A) Changes in the allele frequency of *A*_2_, *D*_2_, and the X- and Y-chromosome in males across 500 generations. (B) Changes in the linkage disequilibrium of *Assister-Distorter, Assister-*XY0, and *Distorter-*XY0 across 500 generations. (C) Haplotype frequencies over 500 generations. *A*_1_*D*_2_X, *A*_1_*D*_2_Y, *A*_2_*D*_2_Y, and *A*_1_*D*_2_0 have near-zero frequencies. The shaded area represents the frequency of *D*_2_ and primarily comprises the haplotypes *A*_2_*D*_2_Y and *A*_2_*D*_2_X. (D) *D*_2_ equilibrium frequency after 1,000 generations. (E) Population sex ratio after 1,000 generations. Parameter values (if not otherwise defined): *k*_*D*_ = 0.25, *s, t* = 0.5, *h*_*A*_ = 0.5, *z* = 0.01, *r* = 0.001, *v* = 0.001.

**Figure 5:**
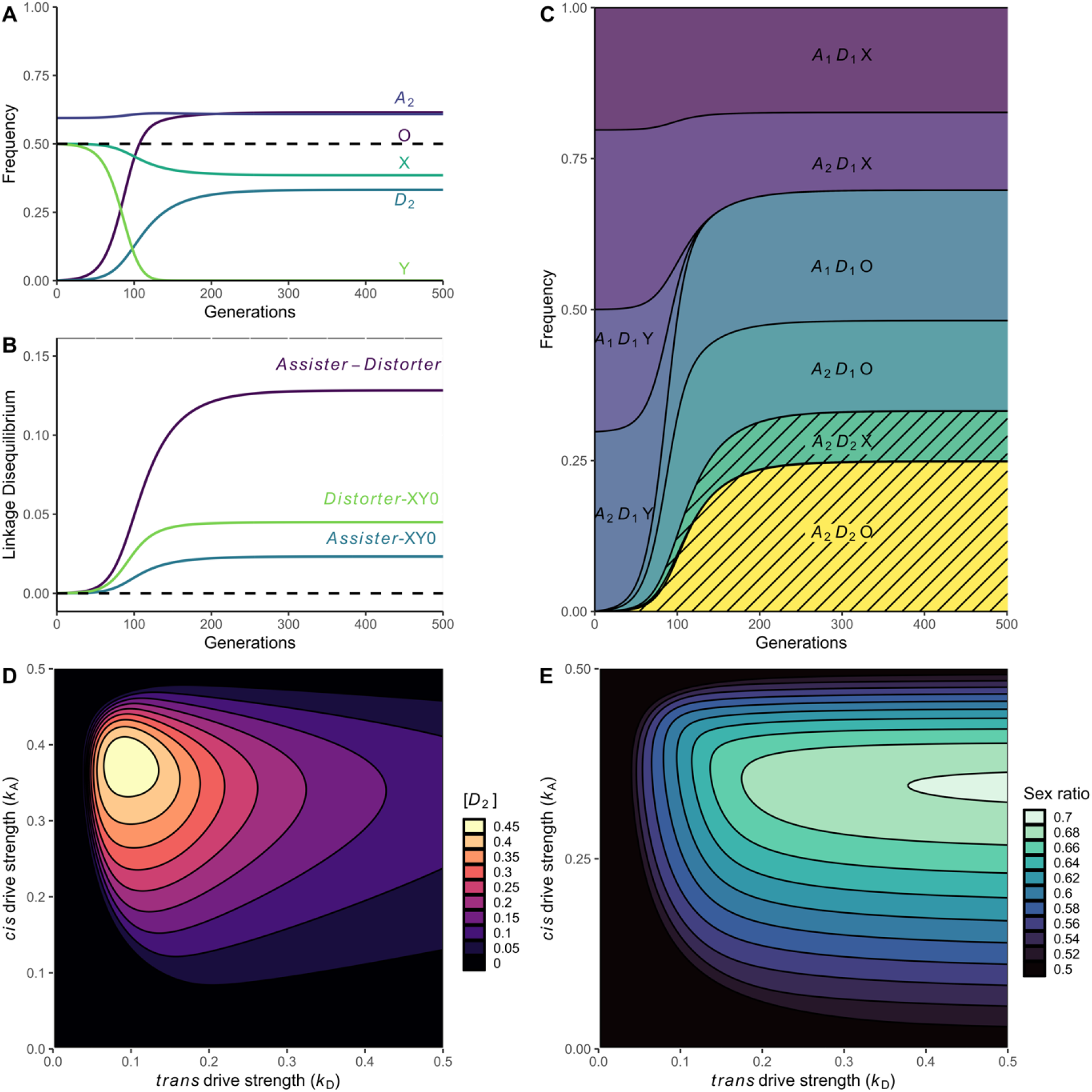
Dynamics and scope of 0-chromosome drive via the *trans*-acting *Distorter* causing X-conversion, linked to a male-limited meiotic driver *Assister*. (A) Changes in the allele frequency of *A*_2_, *D*_2_, and the X-, Y-, and 0-chromosome in males across 500 generations. (B) Changes in the linkage disequilibrium of *Assister-Distorter, Assister-*XY0, and *Distorter-*XY0 across 500 generations. (C) Haplotype frequencies over 500 generations. *A*_1_*D*_2_X, *A*_1_*D*_2_Y, *A*_2_*D*_2_Y, and *A*_1_*D*_2_0 have near-zero frequencies. The shaded area represents the frequency of *D*_2_ and primarily comprises the haplotypes *A*_2_*D*_2_0 and *A*_2_*D*_2_X. (D) *D*_2_ equilibrium frequency after 1,000 generations. (E) Population sex ratio after 1,000 generations. Parameter values (if not otherwise defined): *k*_*D*_ = 0.25, *k*_*A*_ = 0.25, *h*_*A*_ = 0, *z* = 0.01, *r* = 0.001, *v* = 0.001.

## Discussion

Here, we have shown that some *trans*-acting segregation distorters, which target an unlinked locus on the X-chromosome, can spread when they arise on a suitable genetic background; we refer to this novel form of segregation distortion as “remote-control meiotic drive”. Thus, sex-chromosomal segregation distortion need not be caused solely by sex-linked loci. Additionally, our results run counter to the conclusion one may draw from previous theory, which finds that unlinked loci should ordinarily suppress drive (Eshel, 1985; Veller, 2022). We also found that such *trans*-acting distorters could spur a transition in sex chromosome systems from XY to X0 male heterogamety.

We focused on two particular scenarios, where a *trans*-acting segregation distorter (*Distorter*) arises near a locus (*Assister*) that is either under sexually antagonistic selection or exhibits sex-specific *cis*-segregation distortion (in the supplement we consider a third scenario, where the nearby locus experiences parentally antagonistic selection, and a fourth scenario, where the autosomal locus co-segregates with the Y-chromosome). Both genetic backgrounds lead to linkage disequilibrium between *Distorter* and *Assister*, where the *trans*-acting segregation-distorting allele *D*_2_ becomes associated with *A*_2_, the male-beneficial allele or the *cis*-segregation distorting allele in the two respective scenarios. Under either scenario, *A*_2_ has the same optimal destination: a male individual (Figure 6A). If *Assister* is sexually antagonistic, *A*_2_ will act to enhance its bearer’s fitness—and hence its own fitness—when it ends up in a male as opposed to a female. If *Assister* is parentally antagonistic (see Supplementary Material), *A*_2_ will come to be paternally transmitted, and hence will enhance the fitness of its future bearers; this is effectively the same effect as for sexual antagonism, but with the fitness effects delayed by one generation. Lastly, when *Assister* is a sex-specific segregation distorter, *A*_2_ will exhibit a higher-than-Mendelian transmission rate when its bearer is a male, as opposed to a fair transmission rate of 50% in females. In all scenarios, *D*_2_ acts to increase its own effective transmission to males by skewing the sex ratio of its bearer’s offspring. The cost to *D*_2_ of assigning itself to the more common sex is offset by the linked benefit of delivering an *A*_2_ allele to its favored destination (Figure 6B).

**Figure 6:**
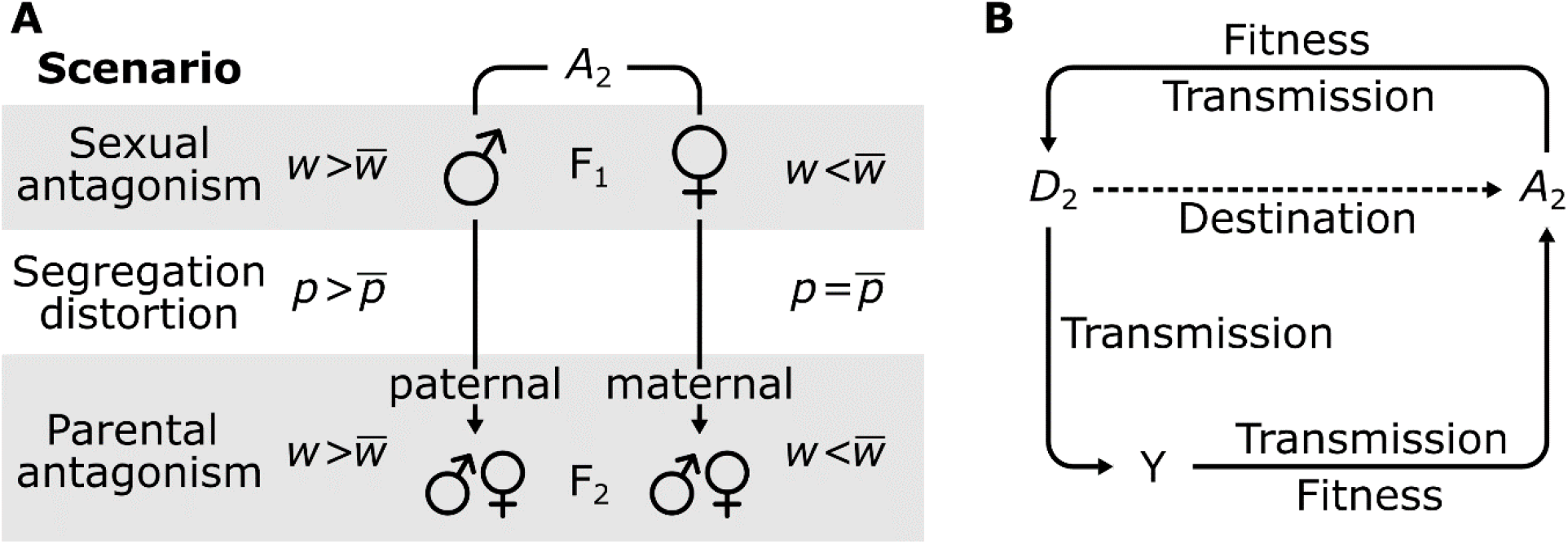
(A) *A*_2_ experiences fitness benefits by preferentially segregating to males under each evolutionary scenario for *Assister*. Sexual antagonism (SA): *A*_2_ confers a fitness benefit so that male bearers have higher-than-average fitness (*w*), whereas female bearers have lower-than-average fitness. Male-limited *cis*-segregation distortion (SD): *A*_2_ has a higher-than-Mendelian transmission (*p*) in males, but regular Mendelian transmission in females. Parental antagonism (PA): *A*_2_ confers a fitness benefit to its bearers when paternally inherited so that they have higher-than-average fitness, whereas it confers a fitness cost when maternally-inherited. (B) *A*_2_ and *D*_2_ each benefit from the other. *A*_2_ provides *D*_2_ a fitness or transmission benefit, and *D*_2_ provides *A*_2_ a “destination benefit,” i.e. indirectly delivers it to a destination where it fares better (i.e., in a male). This synergy is responsible for the gene frequency changes we observe.

The two main scenarios for *Assister* (sexually antagonistic selection *versus cis*-acting drive) have different implications. Empirical studies of these phenomena find that many genes may be under sexually antagonistic selection, but the selective effects of singular loci might be small (Rice, 1992; Innocenti & Morrow, 2010; Schenkel *et al*., 2018). In contrast, segregation distorters are thought to be less common (though many may be cryptic; Werren, 2011), but often involve strong levels of segregation distortion and/or more severe fitness effects (Burt & Trivers, 2006; Lindholm *et al*., 2016). For these reasons, sexually antagonistic selection may enable the evolution of many *Distorter*-like loci throughout the genome, each with either low frequencies of the *trans*-driving alleles or with relatively weak effects thereof. In contrast, segregation distorters might provide fewer opportunities for such loci to evolve, but those that do might harbor stronger or more frequently-occurring *trans*-acting drivers.

Mechanistically, a *trans*-acting sex-chromosome driver may seem rather unlikely. But, on the contrary, it is actually quite easy to obtain. Consider a traditional Y-linked driver that results in biased sex ratios but reaches fixation despite assigning itself to the more common sex. Such sex-ratio biases cause selection to favor the evolution of suppressors on other chromosomes, including the autosomes (Hamilton, 1967; Leigh, 1971; Hurst & Pomiankowski, 1991), to restore the 50:50 sex ratio. This renders the Y-chromosomal driver cryptic, as is commonly seen in e.g. *Drosophila* (Ellison & Bachtrog, 2019; Bachtrog, 2020). Loss-of-function mutations to fixed autosomal suppressors of Y-chromosomal drive should re-enable the initial Y-chromosomal segregation distortion in such mutants. While normally such mutations would be selected against, our model suggests that they may spread if they arise when linked to a locus experiencing balancing selection from either sexually antagonistic selection or sex-specific drive (see also Supplementary Figure S6). In this scenario, the loss-of-function allele at the autosomal suppressor is functionally equivalent to a *trans*-acting distorter of the Y-chromosome. Elsewhere in this paper we have described the *trans*-acting distorter as though it alone caused the drive of what would have otherwise been well behaved, Mendelian sex chromosomes. But in the scenario just outlined the *trans*-acting distorter doesn’t so much cause as *permit* the sex-chromosome drive that had been lying latent. This may also help explain why some suppressors of segregation distortion remain polymorphic rather than become fixed, as is seen in e.g. *Drosophila* (Vaz & Carvalho, 2004; Lyth *et al*., 2023; Gupta & Unckless, 2024).

### Remote-control meiotic drive in relation to sex determination systems

Although we presented a male heterogametic system above, these results might be even more important in female heterogametic (ZW) systems. A key parameter in our models, *z*, measures the fertility effect of sex-chromosome drive. We assume that a male who experiences the loss of some X-bearing sperm is able to compensate those lost sperm with functional Y-bearing sperm, at least to some extent. However, if the fertility effect of drive is sufficiently steep, a *trans*-acting driver may never get off the ground. In ZW species, this sort of compensation is likely automatic. Sex-chromosome drive in ZW systems takes place in an asymmetric meiosis, where some loss of meiotic products (the polar bodies) is a given. Drive here is a matter of navigating to the egg pole during meiosis I rather than sabotaging or incapacitating the half of meiotic products that inherit the alternative allele, as is the case in male meiosis. Because drive in ZW systems may leave fertility unaffected, we might expect *trans*-acting drivers to be more common here.

When remote-control drive occurs through X-conversion, we see that this leads to the evolution of X0 sex determination (females XX, males X0). X0 sex determination is commonly considered to evolve via the loss of the Y-chromosome (Furman *et al*., 2020), owing to its predisposition for accumulating mutations and reduced rates of adaptive evolution (Bachtrog, 2008, 2013). Our model identifies a novel trajectory for this transition that does not rely on the protracted decay of the Y-chromosome, but rather can occur at any point after the original Y-chromosome has evolved. The genetic decay of the Y-chromosome may even make this transition selectively favorable. The “hot potato” model of sex determination transitions posits that the accumulation of deleterious mutations on the Y-chromosome results in a genetic load in males (Blaser *et al*., 2014). This favors males that lack the Y-chromosome, which are normally thought to occur through the evolution of a novel male-determining locus on a (former) autosome. In our model, the conversion of X-bearing sperm into 0-bearing sperm effectively reflects the conversion of a former X-chromosome into a new Y-chromosome (with regard to its male-determining function) (Meisel, 2020). In our analysis, X0 males were assumed to suffer a fitness cost *v* > 0 relative to XY males, but if X0 males actually experience a fitness benefit (i.e. *v* < 0) this should expand the parameter space enabling remote-control drive.

### Remote-control drive has potential practical applications in e.g. pest control

Fundamentally, these results challenge the current view of sex-chromosome drive as a system in which one of a dimorphic pair of sex chromosome skews its own transmission at the expense of another. In our system, the genetic “actors” that encode drive are autosomal, with sex chromosomes involved only for their sex-determining function. This reveals the possibility that the field has previously overlooked cryptic sex chromosome drive systems encoded by autosomal sex ratio distorters and challenges the view that the evolution of Y-chromosome drive is limited by the Y-chromosome’s diminutive size. Moreover, autosomally-encoded sex-chromosome drivers bypass one major obstacle to sex-chromosome meiotic drive: meiotic sex-chromosome inactivation (Meiklejohn & Tao, 2010). This inactivation would prevent X- or Y-chromosomal meiotic drive genes from being expressed during meiosis and is therefore often used to explain the relative dearth of killer sex chromosomes in taxa with meiotic sex chromosome inactivation (MSCI) as opposed to taxa without it (e.g. dipterans) (Burt & Trivers, 2006). However, if the relevant sex-chromosome–meiotic-drive genes are encoded by an autosome rather than a sex chromosome, meiotic sex-chromosome inactivation would not prevent drive.

These observations prompt us to consider the potential application of a *trans*-drive system in the realm of synthetic biology, as autosomally-encoded Y-drives may help overcome current technical barriers. In vector control research, synthetic sex-ratio distortion is an attractive strategy for population suppression of a disease vector such as *Aedes aegytpi*, and researchers have proposed X-shredders to distort population sex ratios, both via autosomally encoded mechanisms (see Alcalay *et al*., 2021) and Y-encoded mechanisms (see Burt & Deredec, 2018). Standard autosomal X-shredders are highly self-limiting (i.e. they only persist for a short period of time), as the autosome encoding drive follows Mendelian inheritance. Autosomal X-shredders therefore require repeated releases of modified mosquitoes to be an effective form of population suppression (Alcalay *et al*., 2021). Y-linked X-shredders are theoretically much more effective than their autosomal counterparts, given their supermendelian inheritance, but are functionally hampered by MSCI (Lifschytz & Lindsley, 1972; Tolosana *et al*., 2024). These shortcomings generate desire for a Y-drive system that both avoids MSCI and is more effective than autosomal drive.

Alongside existing research to circumvent MSCI on the Y-chromosome by inducing expression of the X-shredder before it occurs (Haber *et al*., 2024; Tolosana *et al*., 2024), we suggest additionally considering the potential for remote-control drive as discussed here. Specifically, inserting the synthetic X-shredding mechanism (e.g. CRISPR^SRD^, see Galizi et al. 2016) near either 1) a strongly male-beneficial autosomal allele or 2) a male-limited autosomal driver, could provide a self-limiting synthetic drive more effective than standard autosome-linked X-shredders and yet simpler than Y-linked X-shredders that need to subvert meiotic sex-chromosome inactivation. While ecological modelling is outside of the scope of this paper, we encourage further exploration of the potential application of *trans*-acting drive in synthetic systems.

### Conceptual implications of remote-control drive

Leigh (1971) analogized the genome to a political body, where cabals of a few genes, such as meiotic drivers, would be overwhelmed into fair segregation by a so-called parliament of genes. Specifically, he considered the case of a sex-chromosome driver, whose distortions of the sex ratio would unite any unlinked genes into an alliance to suppress sex-chromosome drive, thus pitting the driving sex chromosome against every other faction of the genome. Ordinarily, this is how selfish genetic elements are viewed: as a small cabal, capable of being put down by the parliament of genes (Scott & West, 2019). But our results call to mind a different political metaphor: politics makes strange bedfellows. We find that unlinked loci may form a cabal, finding a shared interest in skewing the sex ratio. The alignment between autosomal loci and sex chromosomes that we demonstrate here is unexpected on the view that selfish genetic elements must co-locate on what Cosmides and Tooby (1981) called a “coreplicon”. Our findings show that it is not necessarily the transmission pattern that promotes collaboration. Instead, as Gardner and Úbeda (2017) have described, a shared destination (in this case, the male germline), can also enable collaboration among genetic parties.

Finally, the observation that sex chromosomes can theoretically experience *trans*-acting meiotic drive raises definitional and conceptual questions about what meiotic “drive” means. The term “drive,” when first introduced, referred specifically to a gene’s preferential segregation into an egg during female meiosis (Sandler & Novitski, 1957). In the nearly 70 years since, the term “meiotic drive” has taken on a broader meaning, now referring to segregation distorters as well as gamete killers or gamete converters (Burt & Trivers, 2006). Across a broad range of meiotic drive systems, the “driver” is generally taken to mean the element that increases its own transmission and is both the actor and direct beneficiary of drive capabilities. In *trans*-acting drive, however, when the actor (autosome) and the direct beneficiary (Y-chromosome) of drive are physically distinct, it becomes unclear which element is the “driver”. Considering a range of options, if we take the autosome to be the driver, then what is the Y-chromosome? The Y-chromosome could arguably be considered a hitchhiker—though hitchhikers are generally inert rather than an integral part of the system, as the Y-chromosome is here. By contrast, if the Y-chromosome is the driver, then how do we explain its reliance on the autosome? This leads us to our colloquial description of *trans*-acting drive as “remote-control drive,” as this recognizes that the Y-chromosome is the object being driven, while the autosome is the enabler of this system, much like a toy car and its remote.

### Conclusion

Here we have presented a model describing the evolution of a *trans*-acting segregation distorter. Previous theory has assumed that the benefits of segregation distortion must be experienced directly by the genes causing such interference, owing to the costs that come with meiotic meddling. We show that this assumption can be violated; autosomal loci that interfere with the transmission of the sex chromosomes can evolve, provided that they originate in the vicinity of a suitable locus, such as one under sexually antagonistic selection or one that exhibits sex-specific segregation distortion itself. These results show that intragenomic conflict may engender even more exotic forms of selfish genetic elements than previously considered.

## Author contributions

Conceptualization: N.L.G., M.M.P., and M.A.S. Software: N.L.G., M.M.P., and M.A.S. Formal analysis: N.L.G. and M.A.S. Writing – original draft: N.L.G. and M.A.S.. Writing – review & editing: N.L.G., M.M.P., and M.A.S. Visualization: N.L.G. and M.A.S. Supervision: M.M.P. and M.A.S.

## Data availability

The model source code, as well as the data processing, analysis, and visualization scripts are available through https://github.com/MartijnSchenkel/RemoteControlMeioticDrive.

## Acknowledgments

We thank Arvid Ågren for fruitful discussion and helpful comments on an earlier version of the manuscript. The authors declare no conflict of interest.

## Supplementary Material

The results presented in the main text sough to consider the effect of varying the strength of the *Assister* and *Distorter* loci, while keeping other parameters constant and set to optimal conditions (e.g. low fitness cost, low recombination rate). Here, we investigate the effect of varying the fitness cost, *z*, of the *D*_2_ allele, and the recombination rate, *r*, between the *Assister* locus and the *Distorter* locus (Supplementary Figure S1 and S2). In both cases, we find that the sex ratio is most heavily skewed toward males when *z* and *r* are both minimized. The effect of recombination (*r*) seems to be slightly larger than the cost of drive (*z*).

We also present two further selective scenarios for the *Assister* locus, parental antagonism and co-segregation. We consider these scenarios to be less biologically likely than those presented in the main text, though an interesting extension of the same logic that an autosome that favors being in males will enable remote-control Y-chromosome drive.

**Supplementary Figure S1.**
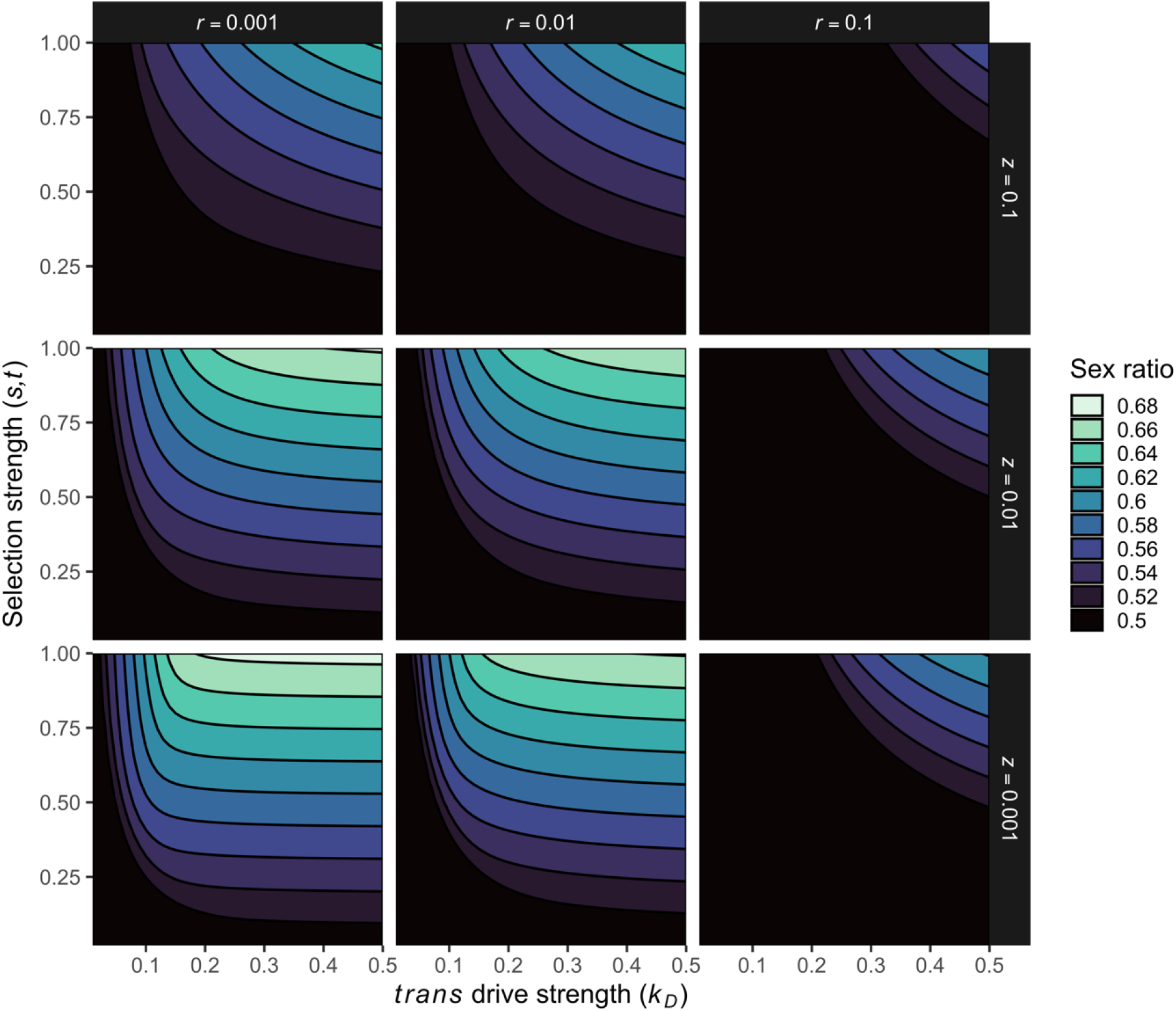
Low recombination rate (*r*) and low *trans*-drive cost (*z*) favor sex ratio distortion when *Assister* is under sexually antagonistic selection. Parameter values: *G* = 1,000, *h*_*A*_ = 0.0.

**Supplementary Figure S2.**
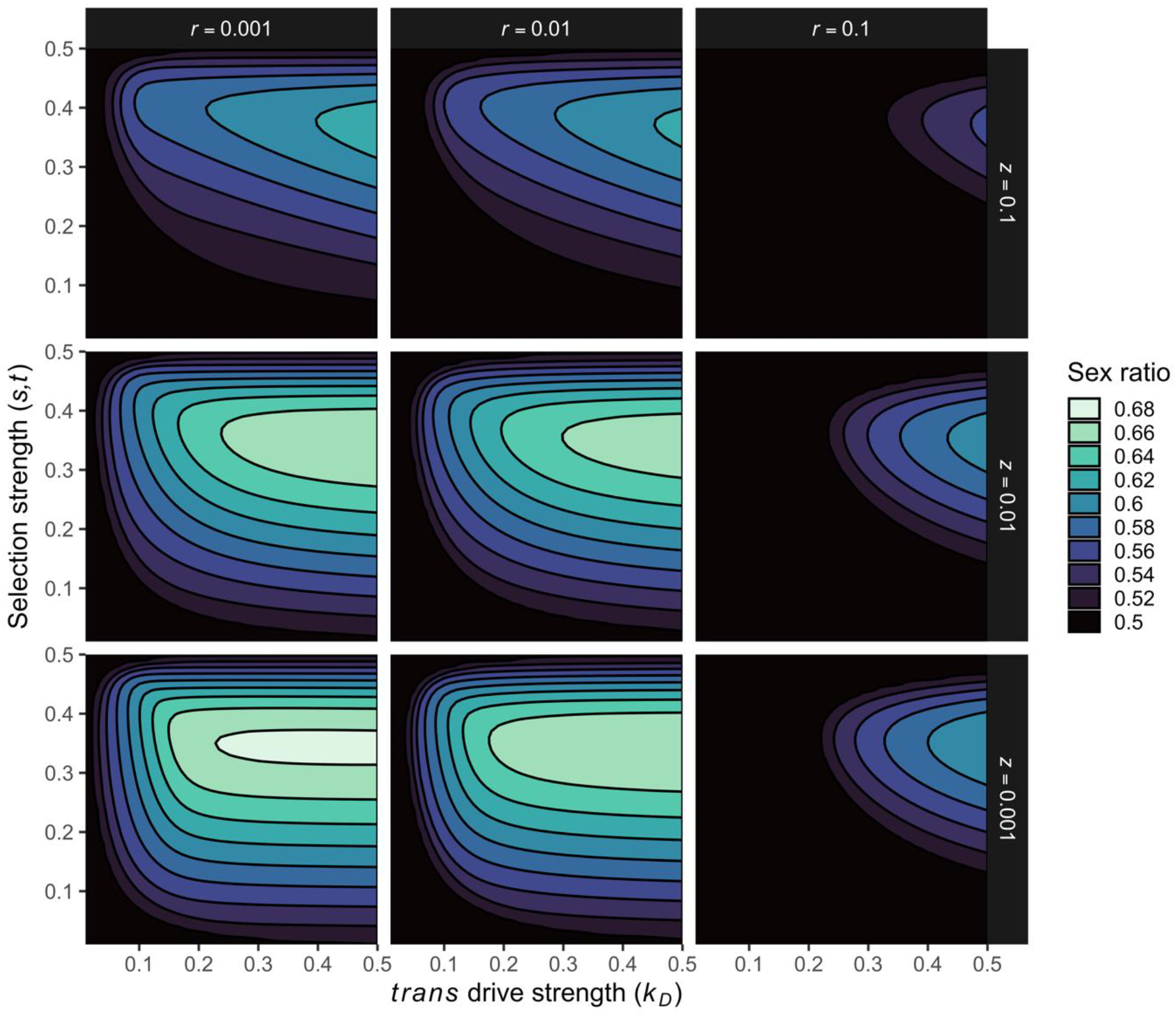
Low recombination rate (*r*) and low *trans*-drive cost (*z*) favor sex ratio distortion when *Assister* exhibits male-limited *cis* segregation distortion. Parameter values: *G* = 1,000, *h*_*A*_ = 0.0.

### Parentally antagonistic selection on *Assister*

Parental antagonism refers to the scenario in which the fitness effect of a gene depends on the parent from which it was inherited. Under parentally antagonistic selection, *A*_1_ and *A*_2_ experience different fitness depending on the parent of origin. We assume that *A*_1_ is beneficial to fitness when inherited maternally and costly when inherited paternally, and vice versa for *A*_2_. The partial fitness score of an individual with genotype *A*_*i*_*A*_*j*_ is given by:

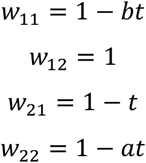

Here, *t* ∈ [0,1] represents the fitness effect of the wrong parent of origin, *a, b* ∈ [0,1] indicates the dominance of a single costly allele when inherited maternally or paternally, respectively.

**Supplementary Figure S3.**
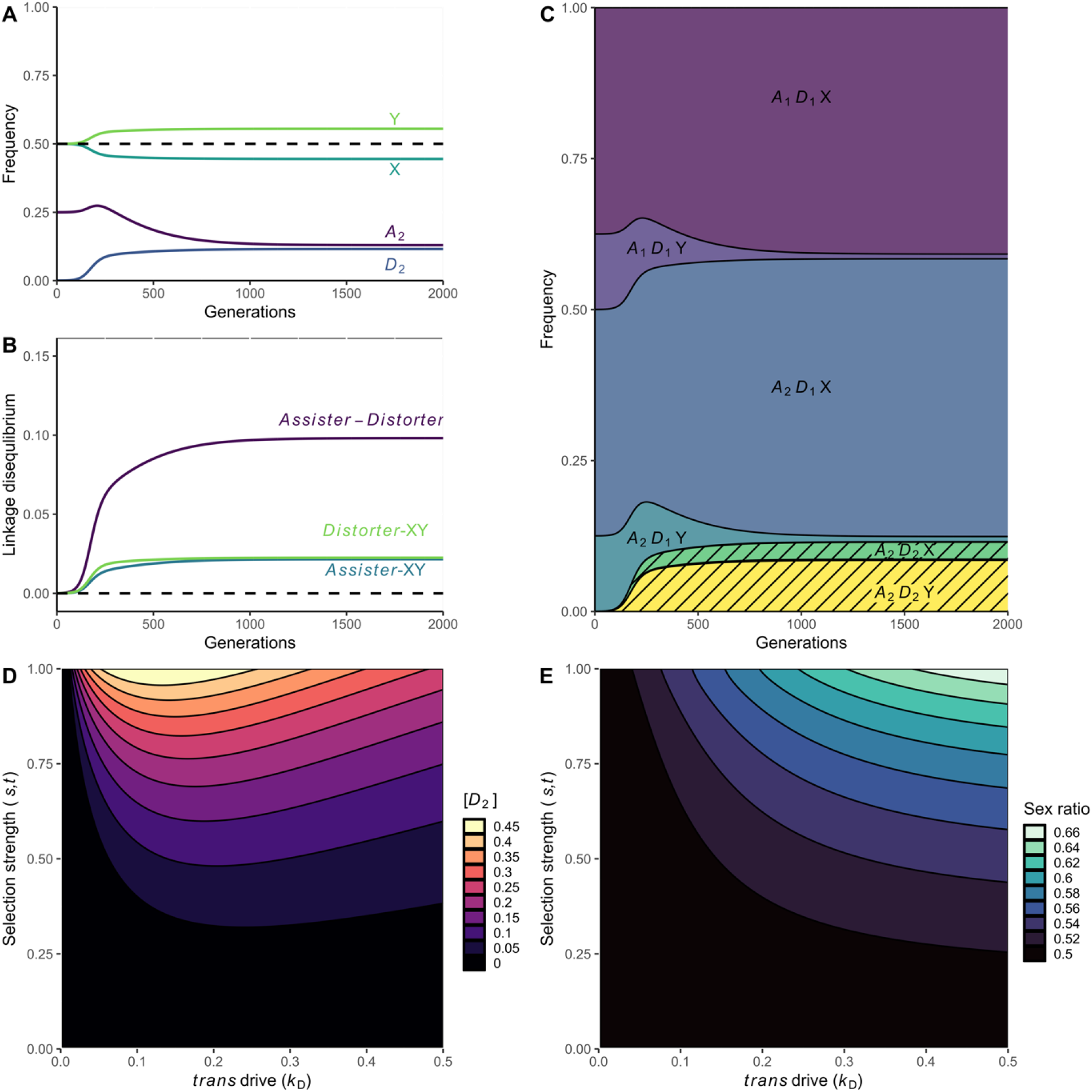
Dynamics and scope of Y-chromosome drive via the *trans*-acting *Distorter* causing X-shredding, linked to a parentally antagonistic *Assister* locus. (A) Changes in the allele frequency of *A*_2_, *D*_2_, and the X- and Y-chromosome across 2,000 generations. (B) Changes in the linkage disequilibrium of *Assister-Distorter, Assister-*XY, and *Distorter-*XY across 2,000 generations. (C) Haplotype frequencies over time. *A*_1_*D*_2_X and *A*_1_*D*_2_Y have near-zero frequencies. The shaded area represents the frequency of *D*_2_ and primarily comprises the haplotypes *A*_2_*D*_2_Y and *A*_2_*D*_2_X. (D) *D*_2_ terminal frequency after 2,000 generations. (E) Population sex ratio after 2,000 generations. Parameter values (if not otherwise defined): *a, b* = 0.5, *t* = 0.3, *k*_*D*_ = 0.25, *z* = 0.01, *r* = 0.001.

### Cosegregation of *Assister*

Our final mechanism for *trans*-acting sex chromosome drive is the most direct: cosegregation of the autosome with the Y chromosome. Here, *A*_2_ encodes a structural variant that creates a physical attachment between the autosome and the Y chromosome. We assume that an *A*_1_*A*_2_ (or *A*_2_*A*_1_) heterozygote, *A*_2_ is co-inherited with Y at a rate of *c* and *A*_2_ with X at a rate of 1 − *c*, where *c* ∈ (0,1] is the cosegregation rate. We assume that *A*_2_ negatively affects its bearer’s fitness, considering that this co-segregation mechanism may induce aneuploidy.

**Supplementary Figure S4.**
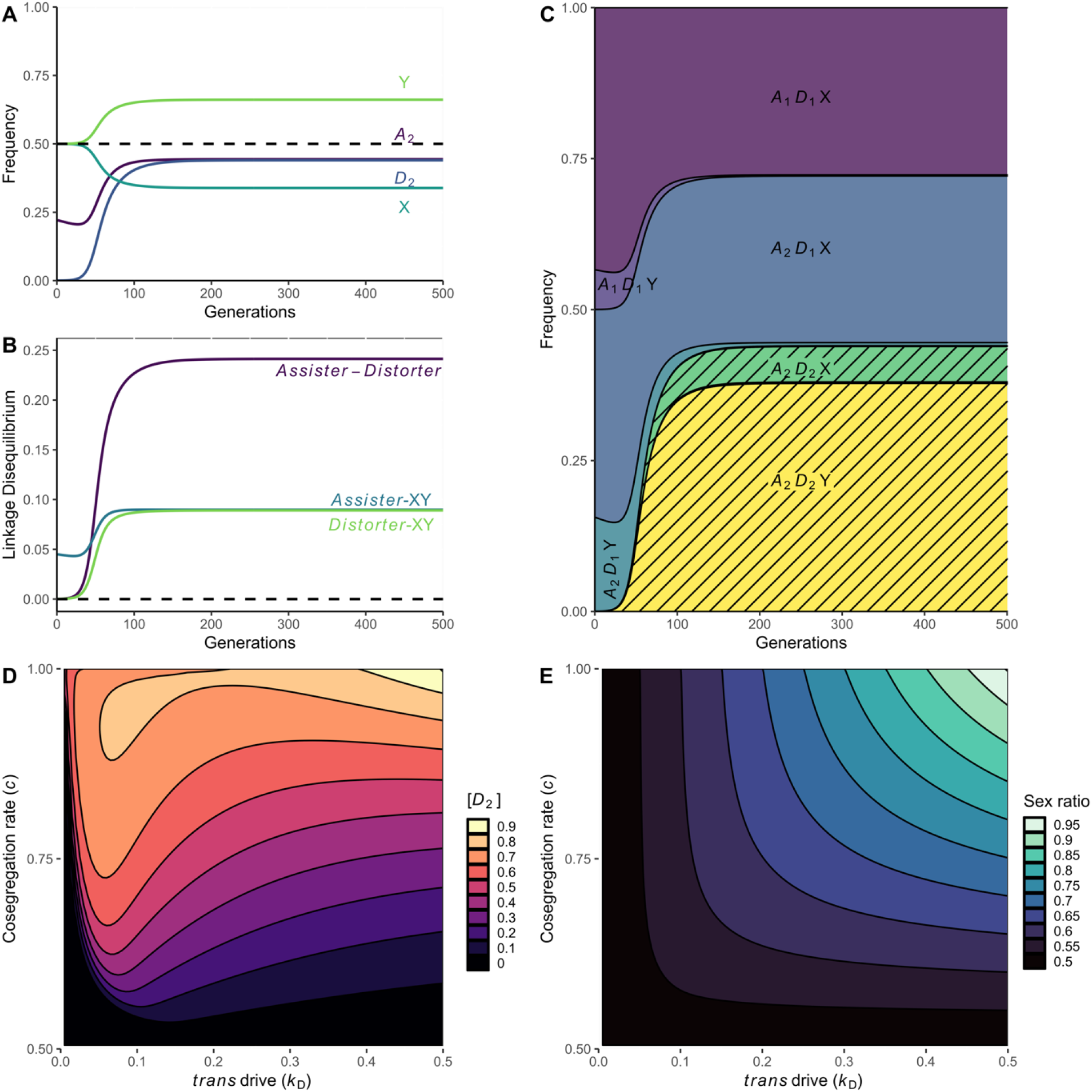
Dynamics and scope of Y-chromosome drive via the *trans*-acting *Distorter* causing X-shredding and exhibiting co-segregation with the Y-chromosome. (A) Changes in the allele frequency of *A*_2_*D*_2_, and the X- and Y-chromosome across 500 generations. (B) Changes in the linkage disequilibrium of *Assister-Distorter, Assister-*XY, and *Distorter-*XY across 500 generations. The shaded area represents the frequency of *D*_2_ and primarily comprises the haplotypes *A*_2_*D*_2_Y and *A*_2_*D*_2_X. (C) *D*_2_ equilibrium frequency after 500 generations. (D) Population sex ratio after 1,000 generations. (E) Population sex ratio after 1,000 generations. Parameter values (if not otherwise defined): *c* = 0.75, *s* = 0.01, *k*_*D*_ = 0.25, *z* = 0.01, *r* = 0.001.

The partial fitness score of a male with genotype *A*_*i*_*A*_*j*_ is given by:

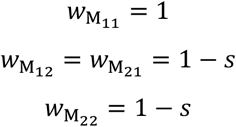

Here, *s* ∈ [0,1] represents the fitness effect of the *A*_2_ allele in males. The partial fitness score of a male with genotype *D*_*i*_ *D*_*j*_ is given by:

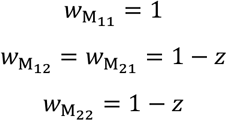

Here, *z* ∈ [0,1] measures the loss of total fertility, and we assume that the *Distorter* locus is neutral in females.

### Polymorphism of XY and X0 sex determination

Considering the XY to X0 transition models presented in the main text, we sought to understand whether polymorphism for the 0- and Y-chromosomes as the male-determining genetic factor was possible. We found a narrow range of possibility, represented in green in Supplementary Figure S5.

**Supplementary Figure S5.**
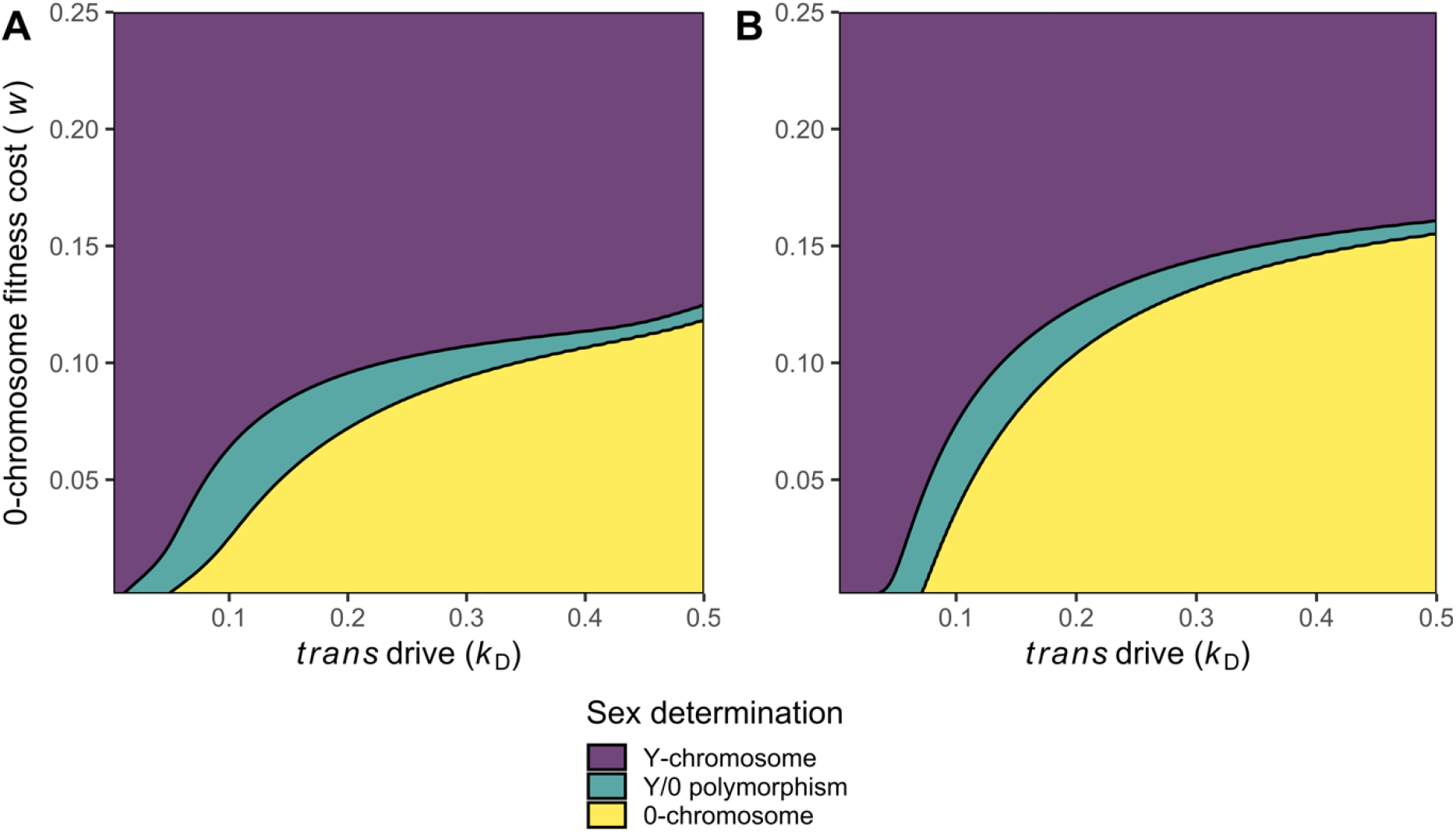
Range of parameter space for which Y-/0-chromosome polymorphism is possible; polymorphism is defined as both the Y- and 0-chromosome having frequencies above 1%. (A) Sexually antagonistic *Assister*, where parameter values: *s, t* = 0.5; *h*_*A*_ = 0.5; *z* = 0.01; ; *r* = 0.001. (B) Male-limited *cis*-segregation distorting *Assister*, where parameter values: *k*_*A*_ = 0.25; *h*_*A*_ = 0.0; *z* = 0.01; *r* = 0.001.

## Supplementary Figures

**Supplementary Figure S6:**
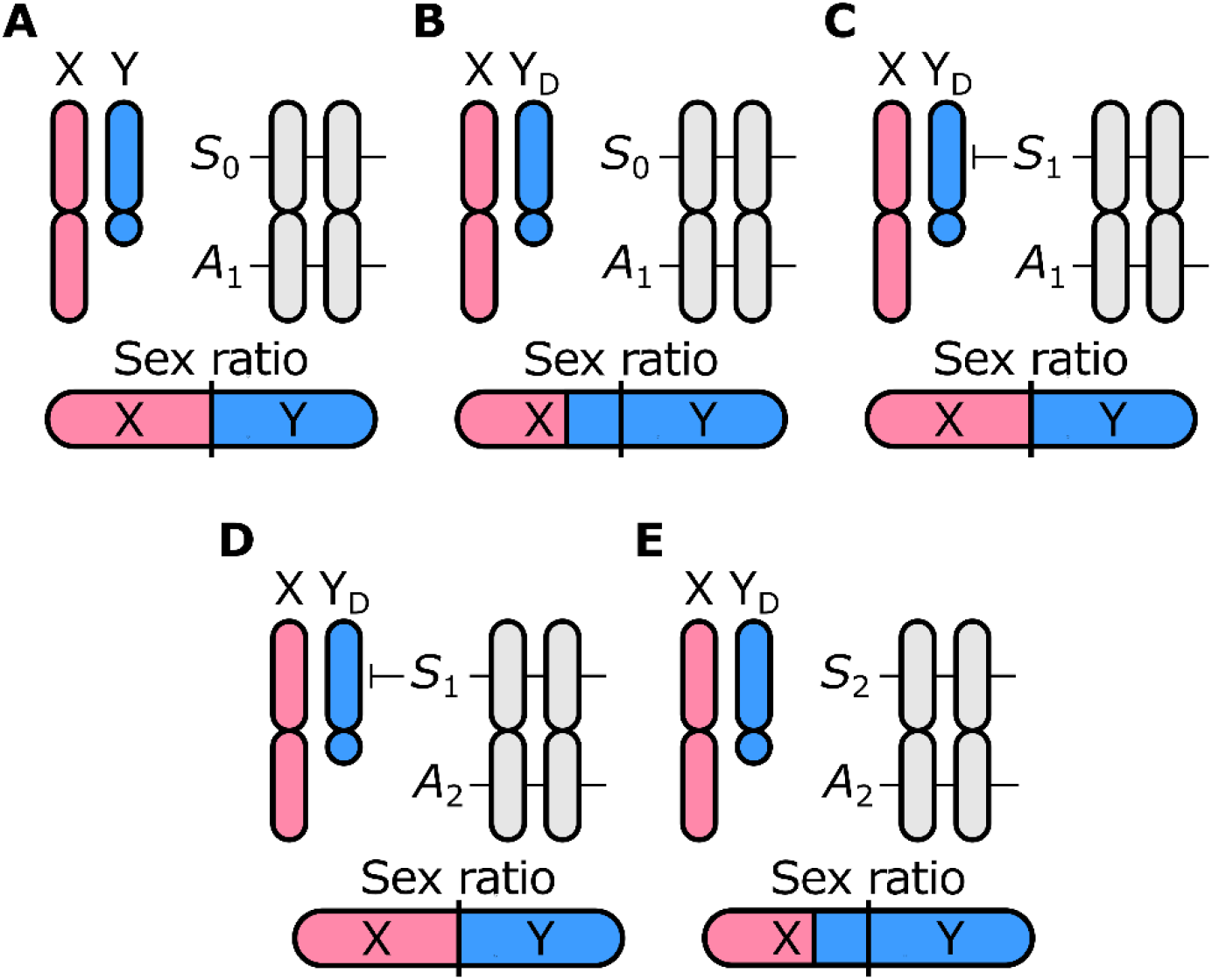
Loss of autosomal suppression of segregation distortion mimics remote control sex chromosome meiotic drive. (A) In the ancestral state, there is no meiotic drive or suppression thereof by any element in the genome. (B) a Y-chromosomal segregation distorter Y_*D*_ evolves and causes male-biased sex ratios. (C) An autosomal locus *S* evolves to reduce Y-chromosomal segregation distortion and restore the sex ratio to equality via the spread of the *S*_1_ allele. (D) A second autosomal locus (*A*), nearby the *S* locus, mutates to yield a novel allele *A*_1_ which exhibits sex-specific segregation distortion. (E) Presence of *A*_1_ renders loss-of-function at the *S* locus advantageous to the *S*-*A* complex; evolution of the non-suppressing *S*_0_ variant re-enables segregation distortion by Y_*D*_, and by extension male-biased sex ratios.

## Supplementary Tables

**Supplementary Table S1:** Overview of model parameters.

| Variable | Description | Standard value or range |
| --- | --- | --- |
| $k_D$ | strength of *trans* segregation distortion | (0,0.5]<br>standard value: 0.25 |
| $z$ | fitness effect of the $D_2$ allele in homozygotes | [0,1]<br>standard value = 0.01 |
| $r$ | Recombination between the *Assister* locus and *Distorter* locus | [0,0.5]<br>standard value = 0.001 |
| $h_A$ | Dominance of $A_2$ | [0,1]<br>standard value = 0.5 for sexual antagonism, 0.0 for *cis* drive |
| $t$ (SA only) | Fitness cost of $A_2$ in females | [0,1]<br>standard value = 0.5 |
| $s$ | SA: Fitness cost of $A_1$ in males<br>*Cis* drive: fitness cost of $A_2$ in both males and females | [0,1]<br>standard value = 0.5 |
| $k_A$ (*cis* drive only) | Strength of *cis* drive | (0,0.5]<br>standard value = 0.25 |
| $b$ | Scalar of less fit paternally inherited allele | (0,1]<br>standard value = 0.5 |
| $a$ | Scalar of less fit maternally inherited allele | (0,1]<br>standard value = 0.5 |
| $t$ | Cost of less fit allele from either parent | (0,1]<br>standard value = 0.3 |
| $v$ | Fitness deficit of X0 males relative to XY males (X-conversion only) | (0,1]<br>standard value = 0.01 |

**Supplementary Table S2:** Overview of fitness scores for all combinations of *Assister* and *Distorter* genotypes.

| Assister type | Assister genotype | Distorter genotype | Female fitness | Male fitness |
| --- | --- | --- | --- | --- |
| Sexual antagonism | $A_1A_1$ | $D_1D_1$ | 1 | $1 - s$ |
| | | $D_1D_2, D_2D_1$ | 1 | $(1 - s)(1 - z)$ |
| | | $D_2D_2$ | 1 | $(1 - s)(1 - z)$ |
| | $A_1A_2, A_2A_1$ | $D_1D_1$ | $1 - h_A t$ | $1 - h_A s$ |
| | | $D_1D_2, D_2D_1$ | $(1 - h_A t)$ | $(1 - h_A s)(1 - z)$ |
| | | $D_2D_2$ | $(1 - h_A t)$ | $(1 - h_A s)(1 - z)$ |
| | $A_2A_2$ | $D_1D_1$ | $1 - t$ | 1 |
| | | $D_1D_2, D_2D_1$ | $(1 - t)$ | $1 - z$ |
| | | $D_2D_2$ | $(1 - t)$ | $1 - z$ |
| Cis-segregation distortion | $A_1A_1$ | $D_1D_1$ | 1 | |
| | | $D_1D_2, D_2D_1$ | 1 | $1 - z$ |
| | | $D_2D_2$ | 1 | $1 - z$ |
| | $A_1A_2, A_2A_1$ | $D_1D_1$ | $(1 - h_A s)$ | |
| | | $D_1D_2, D_2D_1$ | $(1 - h_A s)$ | $(1 - h_A s)(1 - z)$ |
| | | $D_2D_2$ | $(1 - h_A s)$ | $(1 - h_A s)(1 - z)$ |
| | $A_2A_2$ | $D_1D_1$ | $(1 - s)$ | |
| | | $D_1D_2, D_2D_1$ | $(1 - s)$ | $(1 - s)(1 - z)$ |
| | | $D_2D_2$ | $(1 - s)$ | $(1 - s)(1 - z)$ |
| Parental antagonism | $A_1A_1$ | $D_1D_1$ | $(1 - bt)$ | |
| | | $D_1D_2, D_2D_1$ | $1 - bt$ | $(1 - bt)(1 - z)$ |
| | | $D_2D_2$ | $(1 - bt)$ | $(1 - bt)(1 - z)$ |
| | $A_1A_2$ | $D_1D_1$ | 1 | 1 |
| | | $D_1D_2,$ | 1 | $1 - z$ |
| | | $D_2D_1$ | 1 | $1 - z$ |
| | | $D_2D_2$ | 1 | $1 - z$ |
| | $A_2A_1$ | $D_1D_1$ | $(1 - t)$ | |
| | | $D_1D_2,$ | $(1 - t)$ | $(1 - t)(1 - z)$ |
| | | $D_2D_2$ | $(1 - t)$ | $(1 - t)(1 - z)$ |
| | $A_2A_2$ | $D_1D_1$ | $(1 - at)$ | |
| | | $D_1D_2, D_2D_1$ | $(1 - at)$ | $(1 - at)(1 - z)$ |
| | | $D_2D_2$ | $(1 - at)$ | $(1 - at)(1 - z)$ |

## Notes

https://github.com/MartijnSchenkel/RemoteControlMeioticDrive

### Competing Interest Statement

The authors have declared no competing interest.

### Summary of Updates

Figures 2-5 and Supplementary Figures 1-5 have been revised following a slight shift in the model setup. Furthermore, the Introduction and Discussion have been revised to incorporate more extensive information on past work and to elaborate on the scope and interpretation of our results. Lastly, textual edits have been made throughout to e.g. fix typos.

